# SMAD4 Mutation Drives Gut Microbiome Shifts Toward Tumor Progression in Colorectal Cancer

**DOI:** 10.1101/2024.12.20.629749

**Authors:** Travis J Gates, Dechen Wangmo, Kyra M Boorsma Bergerud, Bridget Keel, Christopher Staley, Subbaya Subramanian

**Affiliations:** Department of Pharmacology, University of Minnesota Medical School, Minneapolis, MN; College of Biological Sciences, University of Minnesota, Minneapolis, MN; Department of Surgery, University of Minnesota Medical School, Minneapolis, MN; Masonic Cancer Center, University of Minnesota Medical School, Minneapolis, MN; Center for Immunology, University of Minnesota Medical School, Minneapolis, MN

**Keywords:** Gut microbiome, SMAD4, Driver Gene Mutations, Colorectal Cancer, Metastasis

## Abstract

Colorectal cancer (CRC) progression is driven by a series of sequential mutations in key driver genes, yet the factors underpinning tumor advancement and metastasis remain incompletely understood. Mutations in TP53 and SMAD4, in particular, are associated with poor treatment response and enhanced CRC pathogenesis. Although gut microbiome dysbiosis is implicated in CRC initiation and inflammation, the interactions between the microbiome and specific CRC driver mutations, especially those promoting metastasis, are poorly defined. In this study, we used triple mutant (Apc, Kras, Tp53; AKP) and quadruple mutant (Apc, Kras, Tp53, Smad4; AKPS) organoid-based orthotopic mouse models of CRC to examine the impact of SMAD4 mutation on tumor progression, metastasis, and microbiome composition. AKP and AKPS organoids were endoscopically implanted into the colons of individually housed C57BL/6 mice, and fecal samples were collected weekly over an 8-week period. Our results reveal significant differences in metastatic potential and microbial community dynamics between the two tumor models. AKPS tumors exhibited metastasis to the lymph nodes, liver, and lungs, whereas AKP tumors remained confined to the colon. Longitudinal microbiome analysis showed shifts in microbial composition within each tumor model. Both AKP and AKPS models demonstrated enrichment of *Faecalibaculum* and a decrease in *Dubosiella* over time; however, additional shifts were observed with distinct taxa associated with late-stage tumors in each group. Notably, the AKPS model exhibited higher relative abundances of pro-inflammatory taxa, including *Turicibacter, Romboutsia, and Akkermansia*, suggesting that SMAD4 mutation promotes a more immunosuppressive and pro-metastatic microbiome profile. These findings underscore the role of SMAD4 in modulating the microbiome in a manner that favors CRC metastasis and suggests potential microbial targets for therapeutic intervention to slow CRC progression. This work provides new insights into the microbiome’s role in CRC mutagenesis and metastasis, highlighting the interplay between host genetics and gut microbiota in driving cancer aggressiveness.

## INTRODUCTION

Colorectal cancer (CRC) is the third most diagnosed cancer and the third leading cause of cancer-related deaths in the United States.^1^ Although the global incidence of CRC is declining, early-onset CRC, defined as CRC in individuals under 50, has steadily increased.^1–3^ This trend has led to revised screening guidelines, with colonoscopy now recommended at age 45.^4^ Despite similarities in the genetic mutations found in early-and late-onset CRC, patients with early-onset CRC often have significantly poorer survival rates.^5, 6^

The etiology and oncogenic transformation of CRC involve sequential mutations in key driver genes such as APC, KRAS, TP53, and SMAD4.^7^ Mutations in the tumor suppressor *APC* lead to cellular overgrowth, and polyp formation.^8^ Oncogenic mutations in KRAS promote uncontrolled cell proliferation and resistance to treatment.^9, 10^ TP53 mutations prevent apoptosis in damaged cells, allowing them to proliferate.^11, 12^ The loss of SMAD4 disrupts normal epithelial regulation, promoting excessive differentiation, proliferation, and epithelial-to-mesenchymal transition (EMT), associated with tumor metastasis.^13, 14^

While genetic mutations primarily drive CRC, tumor-extrinsic factors like diet and the gut microbiome significantly influence CRC progression. The diet, characterized by a high fat and sugar intake, has been strongly associated with CRC^15, 16^, with microbiome dysbiosis playing a central role.^17, 18^ Dysbiosis, marked by an increase in pathogenic, cancer-promoting bacteria and a decrease in protective species, can disrupt gut barrier function and foster a pro-inflammatory environment, thus promoting CRC development.^19^ Species such as *Bacteroides* are implicated in this process by compromising the intestinal barrier and facilitating tumorigenesis.^20, 21^

Microbiome alterations are crucial not only in CRC initiation but also in its progression. For example, *Fusobacterium* has been associated with alterations in the ATM gene, which regulates TP53 expression.^22^ KRAS mutations show positive correlations with *Faecalibacterium* and negative correlations with *Bifidobacterium* and *Lachnospiraceae*,^23^ while APC mutations are associated with reduced *Faecalibacterium* and *Bifidobacterium* abundance.^24^ Inactivation of SMAD4 has been linked to decreased *Bacteroides* and increased chemoresistance.^25^

Despite these findings, the longitudinal relationship between the gut microbiome and the accumulation of CRC driver mutations remains underexplored. To address this gap, we used a triple mutant (Apc, Kras, p53; AKP) and quadruple mutant (Apc, Kras, p53, Smad4; AKPS) organoid-based orthotopic CRC model to examine microbiome alterations during tumor progression. Mice with AKPS tumors developed metastatic disease within 8 weeks, while AKP tumors remained localized. This differential tumor behavior prompted us to explore how the presence or absence of SMAD4 might influence the microbial environment and contribute to metastatic burden. Our findings reveal significant microbiome compositional differences between AKP and AKPS mice, with both groups showing increased *Faecalibaculum* as tumors progressed. Conversely, *Dubosiella* was depleted in both groups. Furthermore, *Akkermansia* exhibited inverse trends in abundance, decreasing in AKP mice but increasing in AKPS mice. These results highlight potential microbial factors critical to CRC progression and metastasis.

## MATERIALS AND METHODS

### Organoid generation

C57BL/6 mice harboring germline mutations in *Apc* and *Kras* were used to generate AKP and AKPS organoids for tumor implantation. The colons were carefully dissected and treated with 10 mM EDTA (cat#: AM9260G, Invitrogen, Waltham, MA) to isolate crypt cells, following the protocol described by Sato et al.^26^ The isolated crypts were resuspended in Matrigel (cat#: 354234, Corning, Corning, NY) to support their growth. To create AKP organoids, CRISPR-Cas9 technology was employed to knockout the *p53* gene in the isolated crypt cells. For AKPS organoids, both the *p53* and *Smad4* genes were targeted for knockout using CRISPR-Cas9. The genetically modified crypt cells were cultured in Matrigel, allowing them to form organoids before transplantation.

### Animal husbandry

Female C57BL/6 mice, aged 6 to 8 weeks, were obtained from The Jackson Laboratory. Upon arrival, the mice were individually housed to eliminate potential environmental variations from group housing and ensure consistent conditions throughout the study. The mice were then subjected to an endoscopy-guided injection of AKP/AKPS organoids. Stool samples were collected weekly for the duration of the 8-week study period. At the end of week 8, the mice were euthanized, and their colons were harvested, fixed, and processed for histopathological analysis.

### Endoscopy guided injection

*In vivo* high-resolution colonoscopies were performed using the Mainz Coloview mini-endoscopic system (Karl Storz Endoskope, Tuttlingen, Germany) during orthotopic tumor cell implantation. Mice were anesthetized with 4% isoflurane delivered via inhalation, and atropine (0.04 mg/kg) was administered intraperitoneally to reduce colon motility, contraction, and secretion during the injection procedure. Each AKP group received 5,000 Apc/Kras/p53 cells, while the AKPS group was injected with 5,000 Apc/Kras/p53/SMAD4 cells. Following implantation, the mice were placed on a heating pad to facilitate recovery.

### Longitudinal fecal sample collection

Fecal samples were collected longitudinally once per week for a period of 8 weeks. Mice were transferred to autoclaved, SPF (specific pathogen-free) cages without food or bedding, where they were allowed to remain for one hour to facilitate fecal pellet production. During the collection period, 3 to 5 fecal pellets were harvested from each mouse. Following sample collection, mice were returned to their individual housing. Stool samples were pooled by the experimental group and stored at −80°C until DNA isolation and further processing.

### Histopathological analysis

At the experimental endpoint, mouse colons were fixed overnight in 10% neutral-buffered formalin. After fixation, tissues were transferred to 70% ethanol and submitted to the Clinical and Translational Sciences Institute at the University of Minnesota for further processing. The tissues were paraffin-embedded, and two sections from each mouse were stained with hematoxylin and eosin. Board-certified pathologists subsequently reviewed the stained sections for the presence of dysplasia and both acute and chronic inflammation.

### Fecal DNA extraction and 16S amplicon sequencing

Approximately 0.1 g of mouse fecal pellets were homogenized and processed for DNA extraction using the PowerSoil DNA Isolation Kit (Qiagen) with the QIACube automated platform, following the manufacturer’s Inhibitor Removal Technology (IRT) protocol. The V4 hypervariable regions of the 16S rRNA gene were amplified using the 515F/806R primer set and sequenced. Negative controls, including sterile water and no-template controls, were included to ensure the absence of contamination, and no amplicons were detected. Paired-end sequencing of the amplicons was performed on the MiSeq platform (Illumina, Inc., San Diego, CA, USA), generating 301-nucleotide (nt) reads. All sequence processing was conducted using Mothur software (version 1.48.0).^27^ Raw Fastq files were trimmed to 150 nt to remove low-quality regions and merged using the Fastq-join function.^28^ Sequence reads with mean quality scores < 35 over a 50-nt window, homopolymers > 8 nt, ambiguous bases, or mismatches to primer sequences (more than 2 nt) were discarded. High-quality sequences were aligned to the SILVA database (ver. 132) ^29^ and subjected to a 2% pre-clustering step to remove likely sequencing errors.^30^ Chimeric sequences were identified and removed using the UCHIME package.^31^ Operational Taxonomic Units (OTUs) were clustered at 99% similarity using the complete-linkage algorithm. Taxonomic assignments were made using the Ribosomal Database Project (ver. 16).^32^ The Raw Fastq files are available at https://www.ncbi.nlm.nih.gov/sraPRJNA988535

### Bioinformatics and data analysis

Alpha diversity of the microbial communities was assessed using the Shannon and Chao 1 indices. Beta diversity was evaluated using Principal Coordinate Analysis (PCoA) based on Bray-Curtis dissimilarity matrices.^33, 34^ The Bray-Curtis dissimilarity matrices were computed to measure the pairwise dissimilarity between samples, and PCoA was performed for the ordination of the communities. Differences in beta diversity were further assessed using Analysis of Similarity (ANOSIM),^35^ with Bonferroni corrections applied for multiple comparisons. All statistical analyses were performed at α = 0.05 unless otherwise adjusted for multiple comparisons.

## RESULTS

### Mice with AKPS tumors demonstrated tumor progression and metastasis

We first investigated the differences in tumor progression between AKP (n=9) and AKPS (n=9) mice. Based on the known role of SMAD4 as a regulator of epithelial-to-mesenchymal transition (EMT), we hypothesized that AKPS tumors would exhibit more aggressive behavior than AKP tumors. To test this, we employed an orthotopic injection model in which 5000 organoids carrying mutations in Apc, Kras, and p53 (AKP) or Apc, Kras, p53, and Smad4 (AKPS) were endoscopically implanted into the colon wall of C57BL/6 mice, and tumor progression was monitored over 8 weeks.

Our previous reports and other reports have suggested that AKPS organoids demonstrate metastatic potential in 8-10 weeks.^36, 37^ Interestingly, AKPS tumors showed metastasis to the lymph nodes or liver or both in 6 out of 9 mice, whereas AKP tumors remained localized in the colon of all 9 mice **(Supplemental Fig. S1)**. Histological analysis revealed significant differences in tumor morphology between the AKP and AKPS groups (**Supplemental Fig. S1**). Given the marked differences in metastatic potential and tumor histology, we next examined how the microbiome may be influenced by the presence or absence of the SMAD4 mutation over time.

### Longitudinal changes in the gut microbial community of AKP tumor-bearing mice

We longitudinally examined changes in the microbial community of AKP-tumor-bearing mice (n = 9). Between weeks 2 and 3, we observed an enrichment of *Faecalibaculum* and a concurrent decrease in the relative abundance of *Muribaculaceae* species (**Fig. 1A**). These shifts were accompanied by a temporal reduction in alpha diversity, as measured by the average Shannon index, between weeks 1 and 4 (**Fig. 1B**).

**Figure 1.**
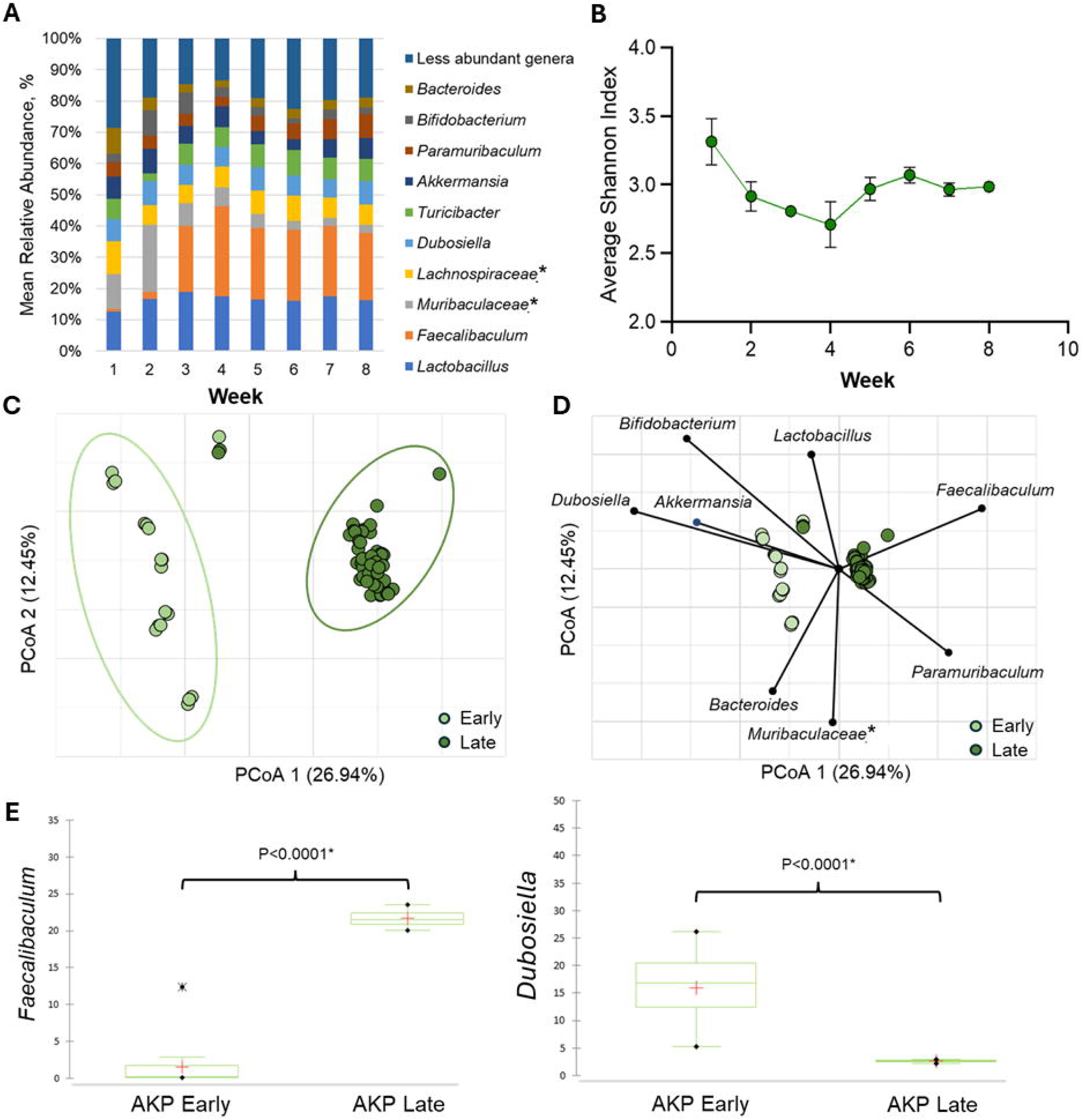
AKP Microbiome Composition and Diversity. **A)** Longitudinal mean percent relative abundance of bacterial genera in AKP tumor bearing mice. Genera that accounted for <0.07 of mean sequence reads were consolidated for clarity. (*) Indicates genera that could not be further classified. **B)** Longitudinal average Shannon index of AKP tumor bearing mice. **C)** Principal Coordinates Analysis of Early (Week 1 – Week 2) and Late (Week 3 – Week 8) microbial compositions. **D)** Correlated taxa with ordination position on either axis (spearman <0.05) are overlaid on the PCoA plot. (*) Indicates genera that could not be further classified. **E)** Kruskal-Wallis comparison of selected bacterial genera between AKP Early (Week 1 – 2) and AKP Late (Week 3 – Week 8). (*p < 0.005)

To investigate overall compositional changes in the microbiome, we performed principal coordinate analysis (PCoA) comparing the early stage (weeks 1-2) and late stage (weeks 3-8) phases of tumor progression. Despite the animals being housed individually, microbiome compositions clustered distinctly by time point, with significant differences between early and late stages, as measured by Bray-Curtis dissimilarity (ANOSIM R = 0.6118, p < 0.001) (**Fig. 1C**).

Next, we explored the correlation between specific bacterial taxa and the PCoA ordination positions to identify taxa associated with early versus late microbiome compositions. Spearman correlation analysis revealed that *Dubosiella* and *Akkermansia* were positively correlated with the early microbiome composition (**Fig. 1D**), while *Faecalibaculum* and *Paramuricaculum* were associated with the late-stage composition.

Finally, we applied the Kruskal-Wallis test to confirm shifts in bacterial taxa between the early and late compositions. We observed significant increases in *Faecalibaculum* (p < 0.0001) and decreases in *Dubosiella* (p < 0.0001) in the late-stage microbiome (**Fig. 1E**).

### Longitudinal analysis of gut microbial community shifts in AKPS tumor-bearing mice

We longitudinally examined microbial community changes in AKPS tumor-bearing mice (n = 9) over 8 weeks. Between weeks 2 and 3, we observed an enrichment of *Faecalibaculum* and a concurrent decrease in the relative abundance of *Dubosiella* (**Fig. 2A**). These microbial shifts were accompanied by temporal changes in alpha diversity, as measured by the Shannon index, a decrease between weeks 1 and 2, followed by an increase between weeks 2 and 6 (**Fig. 2B**).

**Figure 2.**
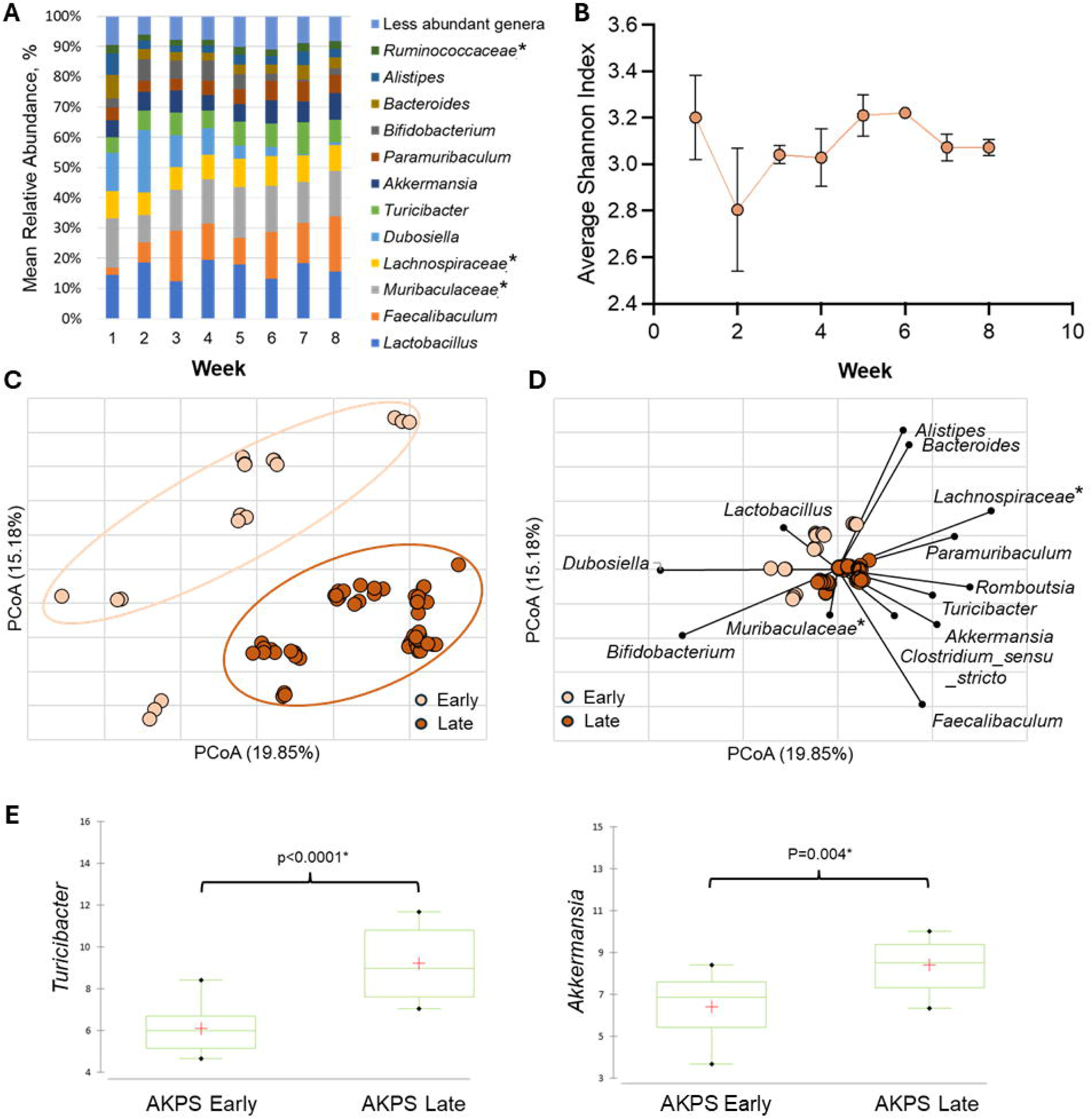
AKPS Microbiome Composition and Diversity. **A)** Longitudinal mean percent relative abundance of bacterial genera in AKPS tumor bearing mice. Genera that accounted for <0.07 of mean sequence reads were consolidated for clarity. (*) Indicates genera that could not be further classified. **B)** Longitudinal average Shannon index of AKPS tumor bearing mice. **C)** Principal Coordinates Analysis of Early (Week 1 – Week 2) and Late (Week 3 – Week 8) microbial compositions. **D)** Correlated taxa with ordination position on either axis (spearman <0.05) are overlaid on the PCoA plot. (*) Indicates genera that could not be further classified. **E)** Kruskal-Wallis comparison of selected bacterial genera between AKPS Early (Week 1 – 2) and AKPS Late (Week 3 – Week 8). (*p < 0.005)

To assess overall compositional shifts, we performed principal coordinate analysis (PCoA), comparing early (weeks 1-2) and late (weeks 3-8) time points. Despite individual housing, significant clustering of microbiome compositions was observed, with a marked difference in microbial profiles between the early and late stages of tumor progression (Bray-Curtis dissimilarity, ANOSIM R = 0.5835, p < 0.001) (**Fig. 2C**).

Next, we identified specific taxa correlated with the early and late microbiome compositions. *Dubosiella* and *Lactobacillus* were compositionally associated with the early stage (**Fig. 2D**), while *Faecalibaculum*, *Turicibacter*, *Romboutsia*, and *Akkermansia* were correlated with the late stage. Kruskal-Wallis tests confirmed significant increases in *Turicibacter* (p < 0.0001) and *Akkermansia* (p < 0.0001) between early and late compositions (**Fig. 2E**).

### Microbial diversity and composition differ between AKP and AKPS tumors

We compared the longitudinal changes in the mean relative abundance of microbial taxa between AKP and AKPS tumor-bearing mice. Both groups exhibited similar trends, with *Faecalibaculum* increasing and *Dubosiella* decreasing as tumors progressed (**Fig. 3A**). To assess differences in microbiome diversity between the two groups, we analyzed alpha diversity. No significant differences were found during weeks 1-2; however, from weeks 3-8, AKPS mice exhibited significantly higher Shannon Index values compared to AKP mice (**Fig. 3B**). There were no significant differences in Chao1 values between the groups (**Supplemental Fig. S2**).

**Figure 3.**
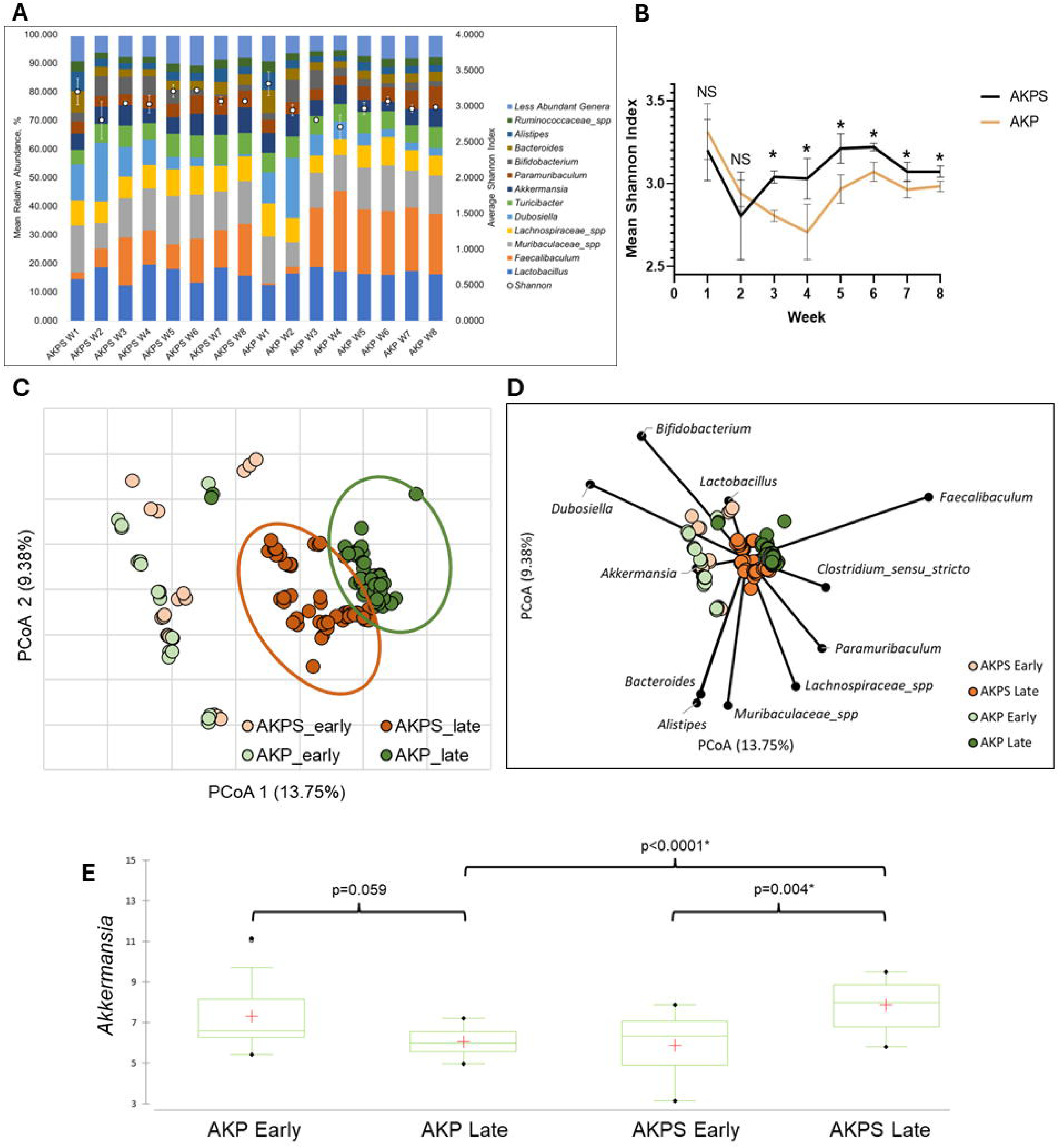
AKP and AKPS Comparative Microbiome Composition and Diversity. **A)** Longitudinal mean percent relative abundance of bacterial genera in AKP and AKPS tumor bearing mice. Genera that accounted for <0.07 of mean sequence reads were consolidated for clarity. (*) Indicates genera that could not be further classified. **B)** Longitudinal average Shannon index of AKPS tumor bearing mice. **C)** Principal Coordinates Analysis of Early (Week 1 – Week 2) and Late (Week 3 – Week 8) microbial compositions. **D)** Correlated taxa with ordination position on either axis (spearman <0.05) are overlaid on the PCoA plot. (*) Indicates genera that could not be further classified. **E)** Kruskal-Wallis comparison of selected bacterial genera between AKPS Early (Week 1 – 2) and AKPS Late (Week 3 – Week 8). (*p < 0.005)

We next assessed differences in community composition between AKP and AKPS tumors using Bray–Curtis dissimilarity (**Fig. 3B**). No significant differences were observed between early AKP and AKPS tumor communities (ANOSIM R = 0.0447, p = 0.125) (**Fig. 3C**). However, the late-stage AKP tumor microbiome composition differed significantly from that of AKPS tumors (ANOSIM R = 0.3725, p < 0.001) (**Fig. 3C**). To identify bacterial genera associated with early and late-stage tumors, we performed Spearman correlation analysis and plotted significant genera on a PCoA (**Fig. 3D**). Early-stage AKP and AKPS tumors were associated with the genera *Dubosiella*, *Akkermansia*, *Bacteroides*, and *Alistipes*. In contrast, late-stage AKP tumors were correlated with *Faecalibaculum*. In contrast, late-stage AKPS tumors showed strong correlations with *Paramuribaculum* and *Muribaculaceae*_*spp*.

To further investigate the differences in bacterial abundance between AKP and AKPS tumors, we performed a Kruskal-Wallis test (**Fig. 3E**). We observed contrasting trends in *Akkermansia* abundance: in AKP tumor-bearing mice, *Akkermansia* decreased between early and late stages (p = 0.059), while in AKPS mice, *Akkermansia* increased over time (p = 0.004). Additionally, *Akkermansia* abundance was significantly higher in mice bearing late-stage AKPS tumors than in late-stage AKP tumors (p < 0.0001). Additional details of significantly altered taxa between early and late stages are provided in **Supplemental Fig. S3**. Finally, we examined longitudinal changes in bacterial genera using the SplinectomeR permuspliner function (**Fig. 4**). We identified significant longitudinal changes in the abundance of several genera, including *Dubosiella* (p = 0.01), *Muribaculaceae_spp* (p = 0.01), *Lachnospiraceae_spp* (p = 0.01), *Turicibacter* (p = 0.03), *Akkermansia* (p = 0.01), *Paramuribaculum* (p = 0.01).

**Figure 4.**
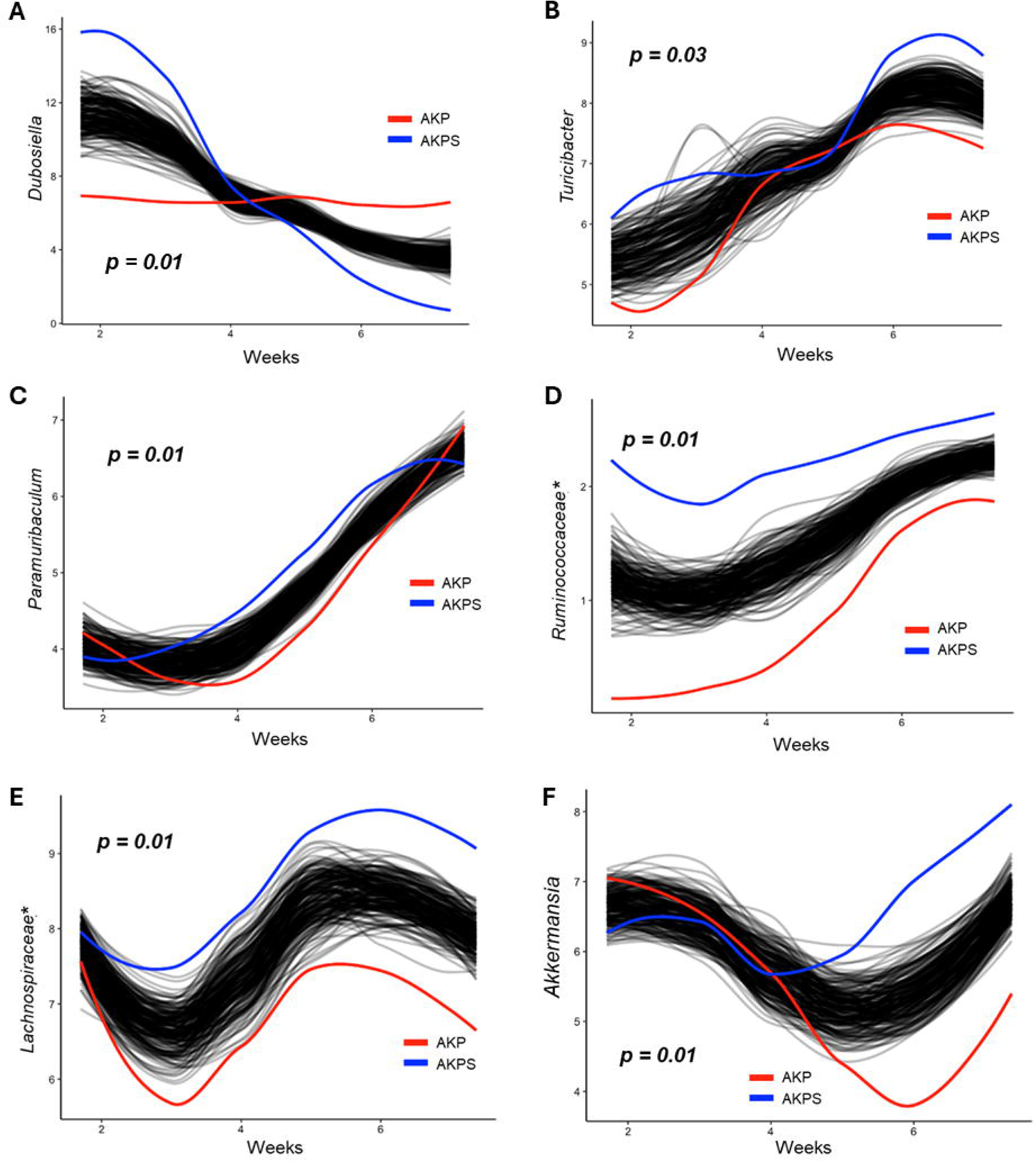
Longitudinally Significant Genera Between AKP and AKPS Tumor Bearing Mice. Longitudinal differences in the relative abundance of significantly different genera between AKP (Red) and AKPS (Blue) tumor bearing mice as determined by SplinectomeR. **A)** *Dubosiella (p = 0.01)* **B)** *Turicibacter (p = 0.03)* **C)** *Paramuribaculum (p = 0.01)* **D)** *Ruminococcacea (p=0.01)* **E)** *Lachnospiracea (p = 0.01)* **F)** *Akkermansia (p=0.01).* (*) Indicates genera that could not be further classified.

## DISCUSSION

Our study provides novel insights into the role of the microbiome in tumor progression and metastasis in colorectal cancer (CRC), focusing on the differential impacts of the SMAD4 mutation. We compared microbial community changes between AKP and AKPS tumor-bearing mice, utilizing a comprehensive longitudinal approach that tracked microbiome shifts over time in relation to tumor progression. Our findings reveal that the microbiome undergoes significant compositional changes in response to SMAD4, providing critical insights into how tumor mutations influence the gut microbial constituents.

Our study revealed that AKPS tumor-bearing mice developed metastases in distant organs, including the lymph nodes, liver, and lungs, while AKP tumors remained restricted to the colon. This finding supports the hypothesis that SMAD4 loss increases tumor aggressiveness, given that SMAD4 is a crucial regulator of epithelial-to-mesenchymal transition (EMT), a critical process for metastasis. Notably, the microbial composition in AKPS tumors differed significantly from that in AKP tumors, especially during later stages of tumor progression, as shown by principal coordinate analysis (PCoA) and Bray-Curtis dissimilarity analysis. These changes suggest that microbial dysbiosis in the AKPS model may contribute to metastatic spread by modulating immune responses or altering the tumor microenvironment to facilitate invasion and colonization of distant organs.

A key finding of our study is the distinct differences in microbial composition between AKP and AKPS tumor-bearing mice. Both groups showed similar trends in *Faecalibaculum* enrichment and *Dubosiella* reduction as tumors progressed. However, longitudinal shifts in microbiome diversity were more pronounced in AKPS tumors, with significantly higher Shannon index values observed between weeks 3-8, suggesting that SMAD4 loss may increase microbial complexity within the TME and potentially influence tumor behavior.

These findings align with emerging evidence linking specific microbial taxa to more aggressive forms of CRC. Although *Faecalibaculum* and *Dubosiella* were key genera in both models, their temporal changes and associations with tumor progression varied between AKP and AKPS mice. *Faecalibaculum*, for instance, correlated more strongly with AKPS tumors in later stages, potentially contributing to immune modulation or promoting an inflammatory environment that favors metastasis. In contrast, *Dubosiella* exhibited a marked decrease in AKPS tumors, indicating its potential as a biomarker for less aggressive tumor types; its depletion may be associated with loss of tumor-suppressive effects or microbial balance. *Dubosiella* has been linked to protective immune suppressive effects in CRC,^38, 39^ suggesting complex interactions between microbial taxa and the immune environment in CRC progression.

The genus *Bacteroides*, known for colonizing mucus and preventing gut dysbiosis by outcompeting pathogens,^40^ showed a declining trend in both AKP and AKPS tumors as they progressed. This aligns with previous findings that link reduced *Bacteroides* abundance to CRC pathogenesis.^41^ *Bacteroides* have been associated with suppressing tumor progression by producing polysaccharide A, which modulates immune responses and maintains gut homeostasis.^42, 43^ Its depletion in AKPS tumors, especially that metastasize, suggests it could serve as a protective agent against tumor advancement and thus may represent a potential therapeutic target.

In contrast, *Bifidobacterium*, typically found at low levels in CRC patients,^44, 45^ increased in abundance over time in both AKP and AKPS tumors. This genus produces lactic acid and acetate— metabolites linked to immune modulation and tumor growth.^46–48^ In colitis, *Bifidobacterium* has been shown to enhance mitochondrial metabolism in regulatory T cells (Tregs).^49^ While Tregs are generally associated with immunosuppression in many cancers, the role of Tregs in CRC has produced mixed results.^50, 51^ The observed increase in *Bifidobacterium* might contribute to an immunosuppressive microenvironment, thereby promoting tumor progression and metastasis.

In AKPS tumors, several genera linked to butyrate and short-chain fatty acid (SCFA) production were more abundant compared to AKP tumors, including *Lachnospiraceae spp.* and *Ruminococcaceae spp.* Notably, *Lachnospiraceae spp.* was present consistently across early to late stages in AKPS samples. Butyrate, produced by these genera, has been shown to modulate inflammatory and immune responses, often promoting an immunosuppressive state.^52–54^ In *APC*-mutated mice, butyrate secretion has also been linked to tumorigenesis.^55^

We also found *Muribaculaceae spp.* to be more abundant in AKPS tumors compared to AKP, although, in AKPS tumors, its abundance decreased from early to late stages. *Muribaculaceae* produces propionate as a metabolic byproduct and has been associated with mucosal repair.^56^ Higher *Muribaculaceae* levels have been inversely correlated with inflammation and carcinogenesis, suggesting its potential protective role in maintaining gut homeostasis and modulating the tumor microenvironment.^57^

The correlation of specific taxa with early-versus late-stage tumor composition reveals important dynamics in the tumor-associated microbiome. For example, *Akkermansia* displayed opposing trends in AKP versus AKPS tumors, with its abundance decreasing in AKP tumors but increasing in AKPS tumors over time. This divergence suggests that *Akkermansia* could serve as a dynamic microbiome marker associated with tumor progression and aggressiveness. The increased presence of *Akkermansia* in late-stage AKPS tumors, particularly in those with metastatic potential, hints at its potential role in promoting metastasis—possibly through modulation of immune checkpoints or alteration of systemic inflammatory responses. *Akkermansia*, a mucin-degrading bacterium, is often elevated in CRC patients.^58, 59^ In our previous studies using the AOM/DSS model of inflammation-driven CRC (id-CRC), we found that fecal microbiota restoration from healthy control mice led to increased *Akkermansia* levels, which corresponded to transient decreases in lipocalin-2, an inflammatory marker in the intestine.^60^ This suggests that *Akkermansia* may have context-dependent roles, influencing tumor progression through immune modulation and interactions with inflammatory pathways.

Additionally, *Lachnospiraceae* spp., known for its butyrate production, was more abundant in AKPS tumors, underscoring the potential immunosuppressive effects associated with SCFA production. Butyrate modulates inflammatory responses in ways that may favor tumor progression.^54^ The enrichment of these microbial genera in AKPS tumors suggests that SMAD4 loss may drive the development of a tumor-supportive microbiome that facilitates tumor progression and metastasis by modulating immune responses and inflammatory pathways.

### Limitations and Future Directions

While our findings are valuable, there are limitations. This study was conducted in preclinical mouse models, and validation in human CRC samples is essential. Our models do not fully replicate the genetic and microbial diversity of human CRC. Future studies will focus on human cohorts to confirm associations between microbial composition and mutations like SMAD4 loss. Additionally, while we observed significant microbial shifts, further research is needed to determine whether these microbes actively influence tumor progression or are bystanders to tumor evolution. Additional studies are required to elucidate the mechanistic role of these microbes in CRC and to explore their potential as biomarkers or therapeutic targets in clinical settings.

### Conclusion

In summary, this study demonstrates that CRC progression is influenced by longitudinal alterations in the microbiome, with distinct shifts occurring in response to the addition of the SMAD4 mutation. These findings provide novel insights into how specific microbial taxa contribute to tumor progression and metastasis, highlighting potential targets for future research.

## Author Contributions

All authors provided meaningful intellectual contributions to this work and were involved in its preparation and editing. TJG, DW, and SS conceived the study. TJG, DW, KMBB, BK, CS, and SS generated and analyzed the data. TJG, KMBB, BK, and SS wrote the manuscript, and all the authors edited it.

## Funding Sources

Research grants from the Mezin Koats Colorectal Cancer Research Fund and Minnesota Colorectal Cancer Funds supported this study.

## ACKNOWLEDGMENTS

We thank the Masonic Cancer Center’s core facilities and the Clinical and Translational Sciences Institute for supporting histological services. Research grants from the Minnesota Colorectal Cancer Funds and Mezin Koat Colorectal Cancer Research Fund support this study.

## CONFLICT OF INTEREST

The authors declare no conflicts of interest.

## ETHICS STATEMENT

This study followed the University of Minnesota Institutional Animal Care and Use Committee protocol code 2107-39273A.

## DATA AVAILABILITY

Raw datasets as Fastq files are available on the NCBI server; https://www.ncbi.nlm.nih.gov/sraPRJNA988535

## SUPPLEMENTAL FIGURE LEGENDS

**Supplemental Figure 1:** Tumor Growth and Morphology Characteristics of AKP and AKPS Tumors:**A)** Table of incidence of primary tumors and metastatic tumor identification in AKP and AKPS tumor bearing animals **B)** Tumor morphology of normal mouse colon, AKP and AKPS primary tumor tissues, AKPS metastatic tumor draining lymph nodes, AKPS metastatic liver tissue

**Supplemental Figure 2:** Chao1 Differences Between AKP & AKPS Mice. Average Chao1 differences between AKP and AKPS Mice. All (p > 0.05)

**Supplemental Figure 3:** Longitudinally Significantly Different taxa Between AKP and AKPS Mice: Kruskal-Wallis comparison of all bacterial genera between AKP and AKPS mice

---

**Graphical Abstract.**
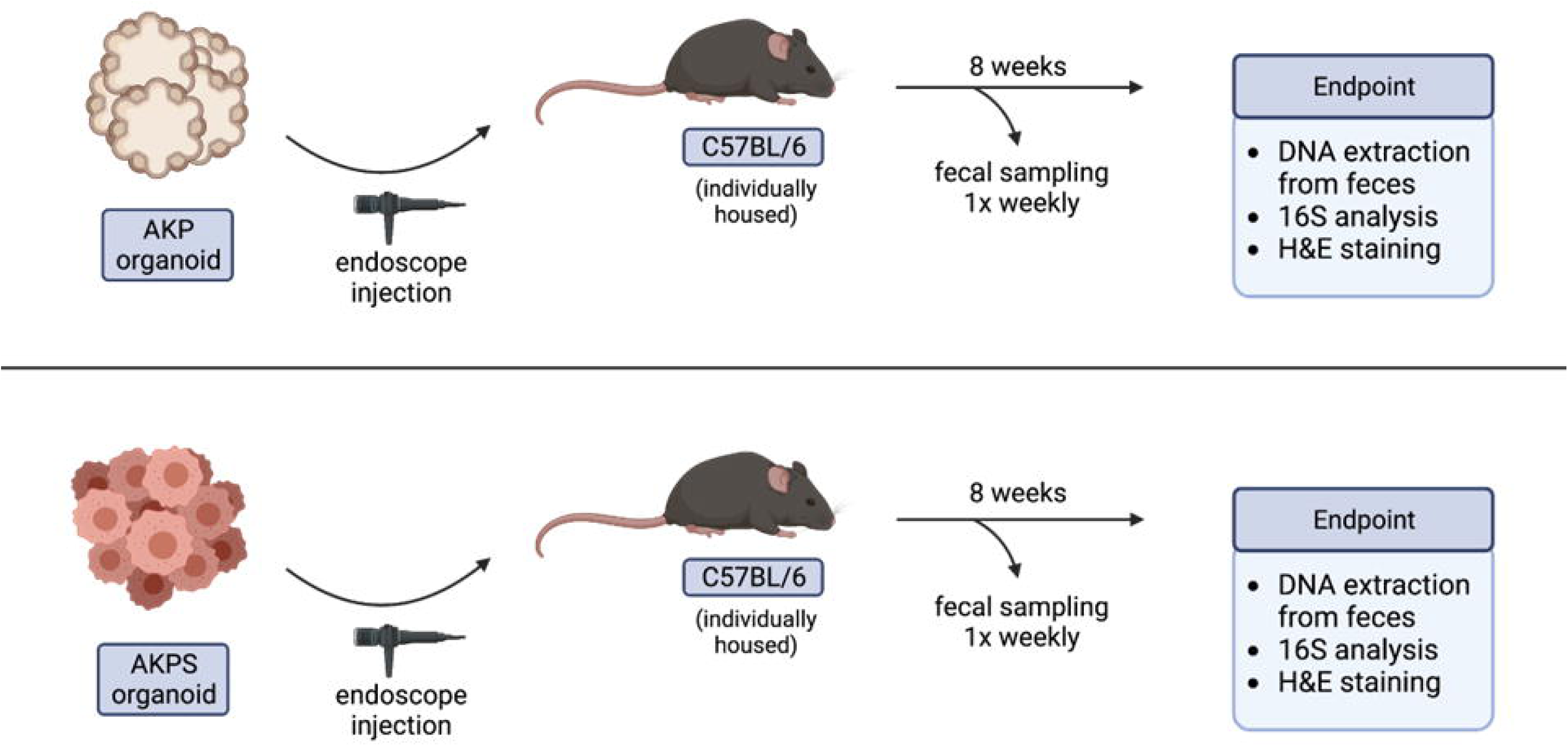

## Notes

### Competing Interest Statement

The authors have declared no competing interest.

