## Supplementary figures and images for "SMAD4 Mutation Drives Gut Microbiome Shifts Toward Tumor Progression in Colorectal Cancer"

### Supplemental Figure 1

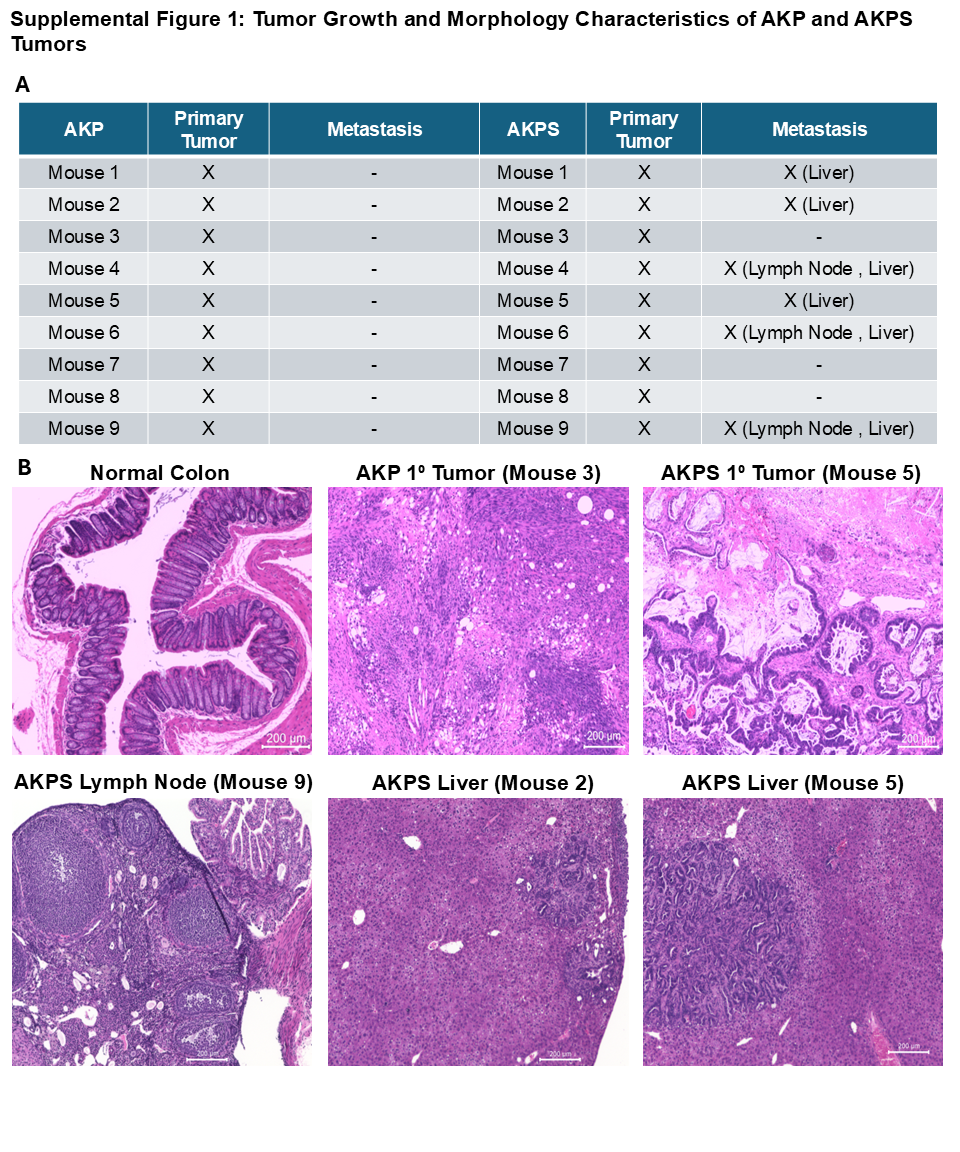

### Supplemental Figure 2

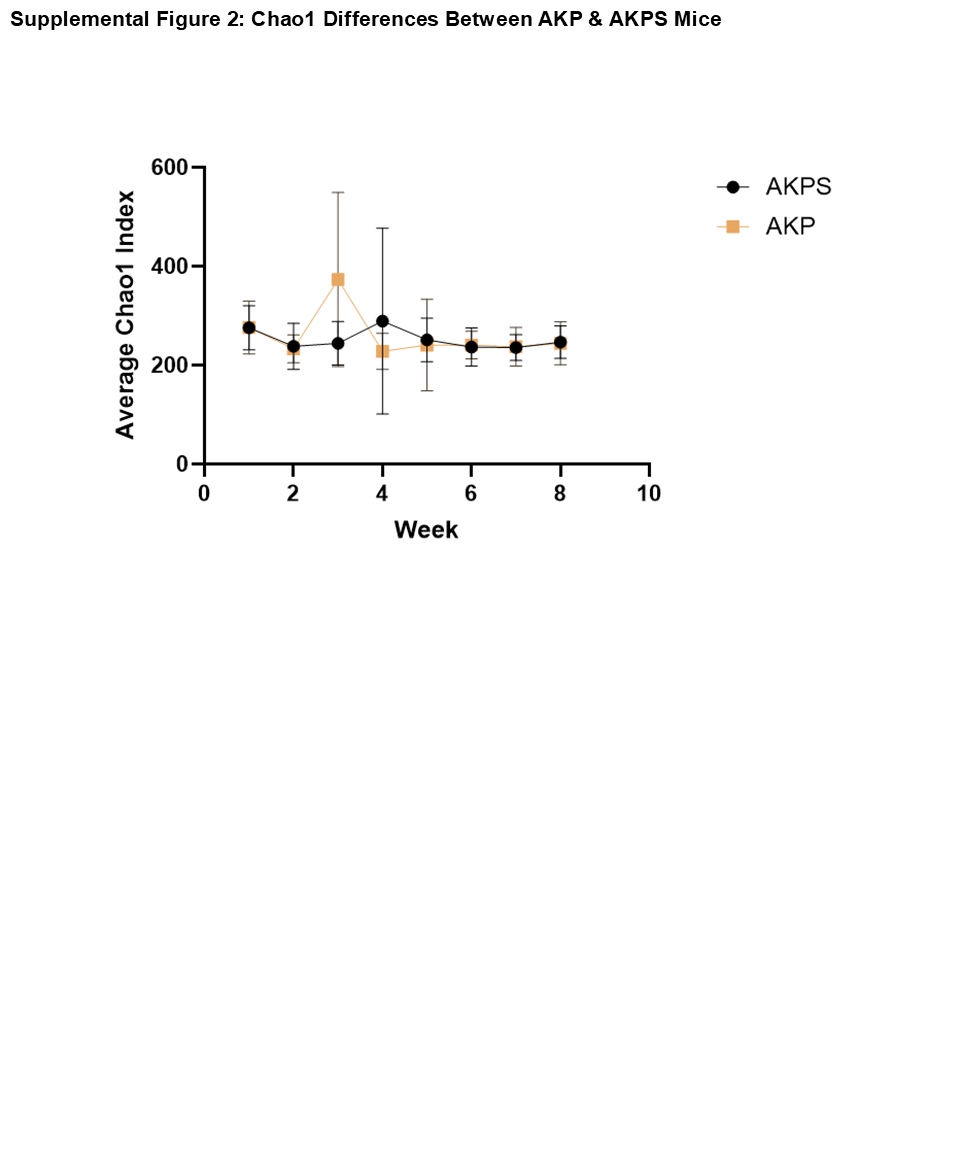

### Supplemental Figure 3

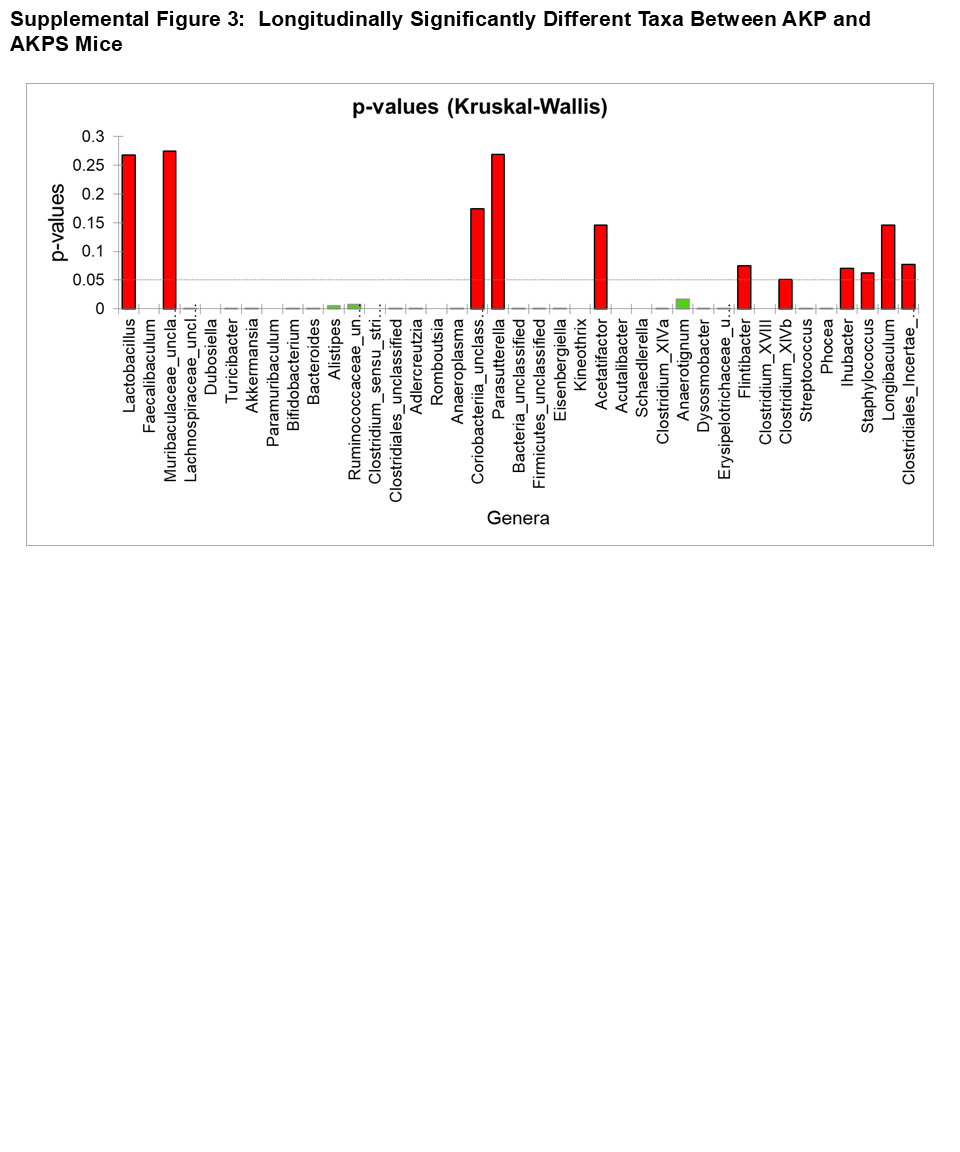
